# Evaluation of domain adaptation approaches for robust classification of heterogeneous biological data sets

**DOI:** 10.1101/682997

**Authors:** Michael Schneider, Lichao Wang, Carsten Marr

## Abstract

Most machine learning algorithms require that training data are identically distributed to ensure effective learning. In biological studies, however, even small variations in the experimental setup can lead to substantial deviations. Domain adaptation offers tools to deal with this problem. It is particularly useful for cases where only a small amount of training data is available in the domain of interest, while a large amount of training data is available in a different, but relevant domain.

We investigated to what extent domain adaptation was able to improve prediction accuracy for complex biological data. To that end, we used simulated data and time-lapse movies of differentiating blood stem cells in different cell cycle stages from multiple experiments and compared three commonly used domain adaptation approaches. *EasyAdapt*, a simple technique of structured pooling of related data sets, was able to improve accuracy when classifying the simulated data and cell cycle stages from microscopic images. Meanwhile, the technique proved robust to the potential negative impact on the classification accuracy that is common in other techniques that build models with heterogeneous data. Despite its implementation simplicity, *EasyAdapt* consistently produced more accurate predictions compared to conventional techniques.

Domain adaptation is therefore able to substantially reduce the amount of work required to create a large amount of annotated training data in the domain of interest necessary whenever the domain changes even a little, which is common not only in biological experiments, but universally exists in almost all data collection routines.

## 1 Introduction

Over the last decade, machine learning, especially supervised learning, has become increasingly important in biological and medical research. Example applications range from protein structure prediction [1,2] and the identification of new disease subgroups from gene expression data [3,4], to the identification of cell connectivity [5] and the prediction of phenotypes from time-lapse [6] data and high throughput imaging [7]. With improving capabilities of data collection and growing computational resources, machine learning will be playing an even more important role in understanding of underlying biological processes.

One of the most well-known limitations of supervised learning, however, is the need for a large amount of annotated data. In biological and medical research, this requirement is often difficult to meet, as it necessitates expert knowledge and intensive manual work. With an increase in high-throughput data it becomes more and more unrealistic to annotate all observations. An appealing alternative is to combine already-annotated data from one or multiple sources in order to build a model for a new problem for which there is only little annotated data.

Another limitation of classic supervised learning techniques is the poor performance in dealing with data from multiple sources. A typical problem in biological research are batch effects. Batch effects describe qualitative changes in measurements because of experimental changes that are unrelated to the biological feature under investigation [8]. Typically, differences in the experimental setup, the use of different protocols, reagents or different machine settings can all lead to such effects. Conventional machine learning techniques are less effective in data with batch effects, due to differences in underlying distributions. Even in the case of an experiment being designed to be a replicate, the classifier trained with data from one experiment often tends to have lower predictive accuracy when applied to data from another replicate [9]. While it is possible to build a new model using only data from one experiment, this would mean wasting expert knowledge and involve labor-intensive annotation for each separate experiment. Consequently, it is desirable to have a model that can achieve a high performance with limited additional annotation work.

Domain adaptation describes the case where at least a part of the data used to train a model follows a different distribution from the data on which the model is finally applied [10]. It is closely related to the notion of transfer learning and mutlitask learning [10,11,12]. We follow Pan and Yang [11] and consider transfer learning as the more general term, with domain adaptation being one special form of transfer learning. Domain adaptation can be applied where a large number of annotated data are available in one or more domains that are not of direct interest (the source domain), while only a limited amount of annotated data is available in the domain of interest (the target domain) (Fig. 1). The idea of domain adaptation is to transfer the knowledge from the source to improve the learning in the target domain. Technically, it can be understood that the pre-trained decision boundary only requires some ‘minor’ tuning from a smaller amout of data to be applied to the new domain. Domain adaptation techniques have originally been developed to address text classification problems [13,14,15].

**Fig. 1.**
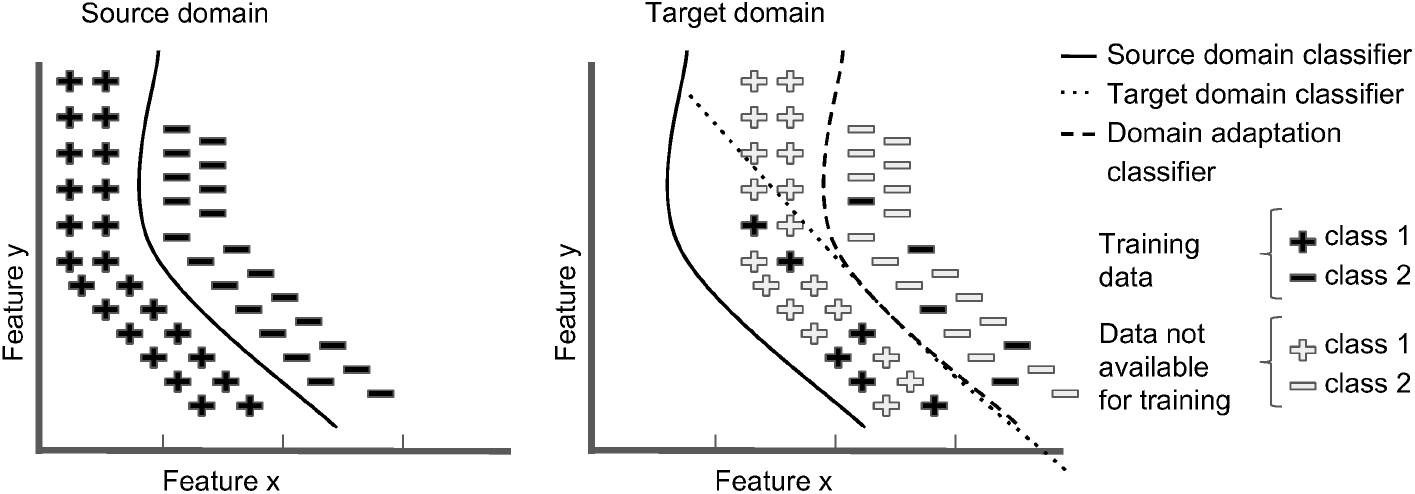
Illustration of a domain adaptation classifier in the target domain that leverages knowledge from a related, but different problem in the source domain. A direct application of the source domain (left) classifier (solid line) would lead to a poor classification in the target domain (right). On the other hand, using only data available in the target domain to train a target domain classifier (dotted line) would also lead to poor performance, as the available data is not sufficient to fully learn the decision boundary. Transferring the knowledge from the source to the target domain using domain adaptation leads to an enhanced classification performance.

Domains in this context correspond to different types, styles or topics, e.g., a model trained with news articles can be adapted to classify a corpus containing fiction texts [14]. However, the concept is very broad and can be applied to any variable that is likely to lead to differences in the data distribution, e.g. different machines, protocols or reagents. Here, we consider domains representing different replicates of a biological experiment, where each replicate can be seen as a different domain.

## 2 Methodology

### 2.1 Definitions

We define a domain *D* as a feature space *X* with the marginal probability distribution *P*(*X*) and a label space *Y*. A function *f*(·) maps *x_i_* to *y_i_*, where *x_i_* ∈ *X* and *y_i_* ∈ *Y*. We consider problems with an arbitrary number of source domains *D*_*s*_1__,…, *D_s_m__* (*m* ≥ 1) and a single target domain *D_t_*. For a multi-class classification problem, we convert to a set of binary classification problems in a one-vs-all manner, i.e. by training a single classifier per class, with the observations of that class as the positive examples and all other observations as negative examples. The aim of domain adaptation is to use the knowledge from the source domains and limited labeling information from the target domain to effectively learn the objective predictive function *f*(·) for the target domain.

### 2.2 Learning techniques

We compare a particular domain adaptation algorithm, the *EasyAdapt* technique [16], with four more conventional techniques of building classifiers. We refer to these as the ‘*Source*’, ‘*Target*’, ‘*Combined*’ and ‘*Domain*’ techniques. In this study, all domains share the same feature space X. In general, the techniques require a common feature subspace across domains. The details of these techniques are outlined below and illustrated in Fig. 2. For all techniques, we assume that the number of observations in the source domains is sufficiently large to estimate a model that will generalize to unseen data from the same distribution. In the *Source* technique, we only use labeled data from the source domains *D*_*s*_1__,…, *D_s_m__* to train the model. The model trained on the source domains is then evaluated on data from the target domain, giving an indirect measure of proximity between source and target domains. In the *Target* technique, we only use labeled data from the target domain *D_t_* to train the model, without considering the data from the source domains. Given enough training data in the target domain, this model should perform the best. In the *Combined* technique, we use labeled data from both the source and the target domains without any reference to the domain membership when training the models (where every data point is weighted equally). This is arguably one of the most common approaches in practice [17,18,19], where a typical scenario consists of a relatively large amount of labeled data from the source domains and a limited amount of data from the target domain. In the *Domain* technique, we slightly adapt the *Combined* approach. An additional set of binary variables encoding the domain membership, in the form of one-hot-encoding, is added to the existing feature set [20]. It is expected to enable the estimated function to have a different offset for each domain, while making use of all the other predictors from all domains to define the shape of the function in common. The *EasyAdapt* domain adaptation technique [16,21], uses a simple transformation to create a representation for the general data structure common to source and target domains and a separate representation for each domain. The transformations 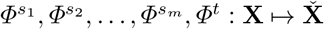 between the features spaces of the different domains have the following form:

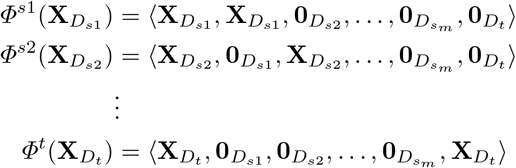

0*_D_d__* denotes a matrix of dimensions corresponding to the dimensions of domain *d* filled with zeros. *EasyAdapt* can be applied to an arbitrary number m of source domains *D*_*s*_1__,…, *D_s_m__* and a single target domain *D_t_* (see Fig. 2 for a visualization and a comparison with other techniques). Features only available in the target domain could also be incorporated by setting the relevant entries for the other domains to 0. The technique is simple and flexible and can be used with any supervised classifier. However, it is recommended that the number of features per domain is not too large, because the feature space increases to ℝ^(*m*+2)*p*^ dimensions with *p* being the dimension of the shared feature space.

**Fig. 2.**
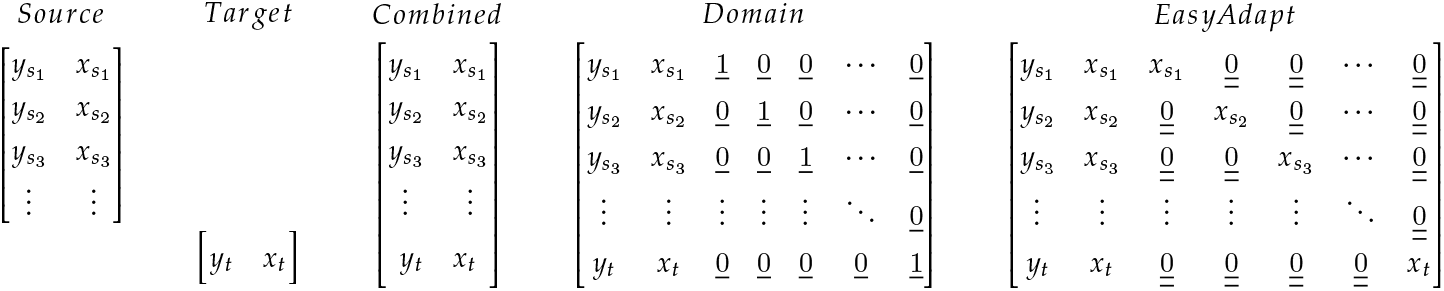
Schematic overview over the different learning techniques. We denote the feature matrices with *x*_*s*_1__ to *x_s_m__* for the *m* source domains and with *x_t_* for the target domain. Label vectors are denoted by *y_s_i__* and *y_t_*, respectively. Single underlined zeros and ones are column vectors, while double underline indicates matrices of dimensions matching the dimensions of *x_i_*. The *Domain* technique is adding an additional feature encoding the domain membership in the form of a one-hot encoding, where the *k*th domain is encoded via a 1 at position k. The *EasyAdapt* technique creates both a unified representation of the data across all domains (analogously to the *Combined* technique) and a separate representation for each domain (diagonal entries).

## 3 Results

### 3.1 Simulation study

In order to visualize how the different techniques work and to test their performance, we created a two dimensional artificial data set with one source domain and one target domain (each with 200 data points), where the ground truth is known (see Fig. 3A). The data was created as follows: In the source domain, we simulate the positive class by sampling 200 data points uniformly around a central point with coordinates (1.0, 0.0). The distance from the centre is sampled from a uniform distribution with mean 0.5 and a range between 0.1 and 0.9. The radial angle is uniformly distributed between 0 and 360 degrees. For the negative class, 200 data points are sampled uniformly around the same central point, but the distance from the centre is sampled from a uniform distribution with mean 0.9 and a range between 0.5 and 1.3. Again, the radial angle is uniformly distributed between 0 and 360 degrees. In order to create the data for the target domain, we translate both classes in the source domain by *y*′ = *y* − 0.60, where *y* is the horizontal coordinate in the source domain while *y*′ is the horizontal coordinate in the target domain. 15% of the data in the target domain was used for training. The remainder of data in the target domain was used for performance evaluation. Support Vector Machine (SVM) [22,23] with a radial basis function (RBF) kernel was chosen as the basic classifier for all the five learning techniques described in the previous section. Parameters were selected using a grid search with 5-fold cross-validation. From both the contour lines (Fig. 3B-F) and the ROC curves (Fig. 3G) it is evident that the *EasyAdapt* technique captured the distribution of the target domain most accurately (AUC = 0.91), by leveraging information from both the source domain and the limited amount of training data from the target domain in building the classifier. Fig. 3B illustrates that due to the limited amount of training data in the target domain, the *Target* technique (AUC = 0.86) learned a decision boundary that was much more complicated than the underlying distribution. The *Source* technique (AUC = 0.55, Fig. 3C) directly applied the decision boundary learned from the source to the target domain, leading to an evident discrepancy with respect to the target domain distribution. The *Combined* technique (AUC = 0.64, Fig. 3D), shifts towards the target domain when building the model. Due to the comparatively large number of source domain data, however, the model is strongly biased towards the source distribution. The *Domain* technique (AUC = 0.89, Fig. 3E) learned a model that describes the target domain quite well, especially in regions close to the centre. In regions that were farther away, however, the contour lines were clearly distracted by source domain information. Compared with these four techniques, the *EasyAdapt* technique (Fig. 3F) learned a model that described the target distribution the best, by successfully integrating the information from the two domains.

**Fig. 3.**
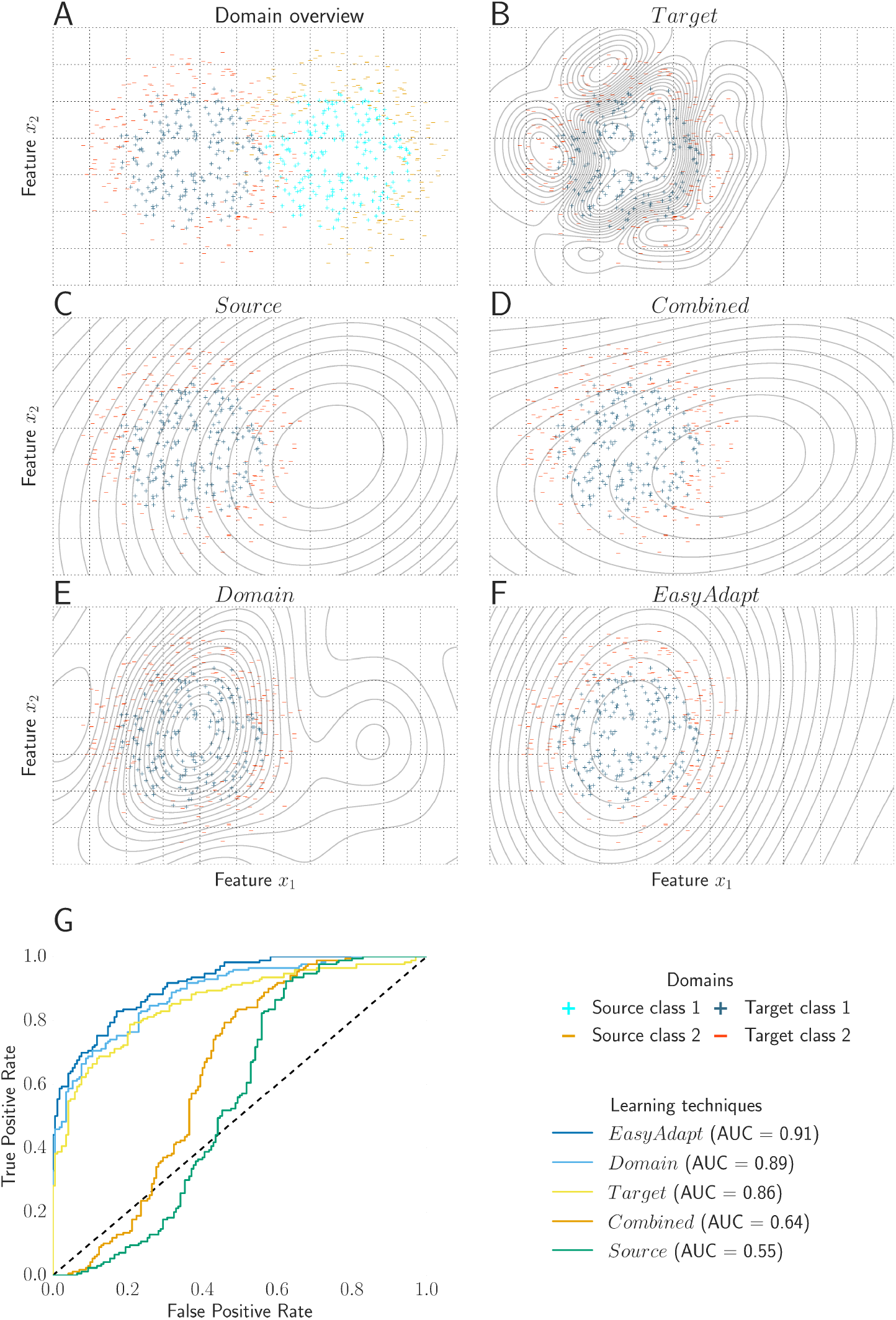
Simulated data: with limited training data and sufficient domain similarity, *EasyAdapt* has the best classification performance on the target domain. (A) Distribution of the two classes in the source (light blue and orange symbols, right) and target domain (blue and red symbols, left). The target domain was divided into a training set and a test set. The target training set consisted of 15% randomly sampled data from the target domain. Classifiers were trained using RBF kernel SVM. (B-F) Classifiers created using *Target* (B), *Source* (C), *Combined* (D), *Domain* (E) and *EasyAdapt* (F). Contour lines represent different thresholds of the decision boundary of the corresponding classifier. (G) ROC curves for the different techniques.

### 3.2 Imaging data set

For a realistic evaluation case, we applied the techniques to a biological data set [25] consisting of 2888 cells with 186 cell texture and shape features from time lapse microscopy experiments, where 8 different cell cycle stages have been manually annotated. The data comes from three experiments, with 1468, 726, and 694 cells, respectively. It is important to note that the experiments differ regarding the microscope objectives and the magnification factor (10x for experiments 1 and 3, and 20x for experiment 2) used, and were conducted by different lab technicians [25]. The different techniques were trained and tested in a one-vs-all manner on the 8 cell cycle stages (where each stage is treated as a separate class). We always picked two experiments to represent the source domains and the remaining experiment as the target domain. We tested all three possible combinations of two source domains and one target domain. All data from the source domains together with the data from the target train set were centered and scaled to unit variance. Subsequently, we applied a principal component analysis (PCA) to the data, (i) keeping only factors explaining 98% of variance (reducing the number of features to roughly 20-30), and (ii) keeping only the 16 highest loaded principal components. We used 4-fold cross-validation and a grid search to select parameters and subsequently evaluated performance on a test set in the target domain. The procedure was repeated 50 times for different target training set sizes of 100, 120, 150, 200, 250, 300, and 400 samples in order to obtain robust estimates for variable performance, especially when using small training set sizes. Independent of the amount of data available in the target domain, we used a fixed-sized test set with 240 samples for performance evaluation, which was randomly chosen for every iteration and for every new training set. In order to evaluate and compare performance of techniques, we chose the microaveraged AUC. Using this metric, class imbalances were taken into account by computing cumulative values for true positives, false negatives, true negatives and false positives for every label and then computing the performance measure from the aggregated values [24]. We compared three different base classifiers, namely a linear SVM [23], an RBF kernel SVM [22], and a random forest classifier [26].

We found that the *EasyAdapt* technique is particularly robust when working with a small set of training samples in the target domain and consistently performed among the top techniques in the regime of small training set sizes (Fig. 4). As expected, with increasing training set size the *Target* technique catches up and for 400 training samples (the maximum training set size in the study), the performance for this technique was among the best performing techniques. In general performance improved for all techniques with increasing training set size with exception of the *Source* technique, which was not trained with any of the target domain data. Results from all experiments are summarised in Table 1, showing the performances of the five learning techniques across three different base classifiers, two different feature selection methods and three different target domains (each combination of a base classifier, a feature selection method and a target domain is referred to as a ‘setting’ below).

**Fig. 4.**
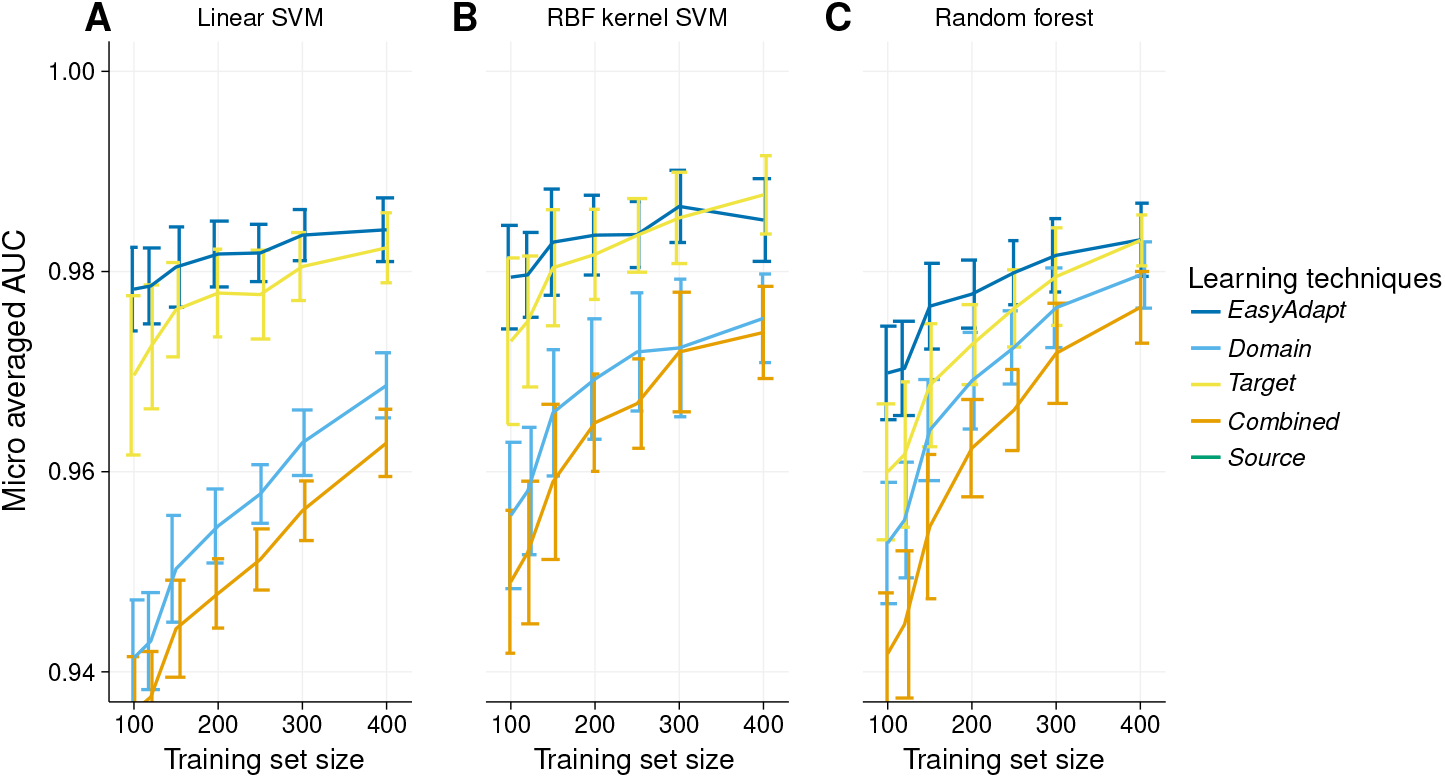
*EasyAdapt* outperforms other techniques in particular for small training set sizes. Performance for (A) linear SVM, (B) radial basis function (RBF) kernel SVM, and (C) random forest classifiers for learning with experiments 1 and 3 as source domains and experiment 2 as the target domain. Performance is measured as microaveraged AUC (mean±standard deviation, n=50 iterations) [24]. We do not plot the *Source* technique since it is independent of the training set size.

**Table 1.** Mean micro-averaged AUC for different classification methods, learning techniques, feature sets (see text for explanation), and target domains. The best performing technique in a row is marked in bold. Note that the performance is averaged over the full range of training set sizes and that one value in the table corresponds to an average of the performance across different training set sizes. Thus, while the average performance for the *Target* technique appears relatively high, it is much lower when the target training size is small. *EasyAdapt*, on the other hand, consistently outperforms other methods, when the target training data size is small (e.g., 100 – 200 instances, see Fig. 4).

| Method | number of features | Target domain | *EasyAdapt* | *Domain* technique | *Target* technique | *Combined* technique | *Source* technique |
| --- | --- | --- | --- | --- | --- | --- | --- |
| linear SVM | 16 | 1 | **0.972** | 0.970 | 0.971 | 0.959 | 0.952 |
|  |  | 2 | **0.978** | 0.949 | 0.976 | 0.941 | 0.803 |
|  |  | 3 | **0.987** | **0.987** | 0.986 | 0.983 | 0.981 |
|  | 98% | 1 | **0.976** | 0.974 | 0.974 | 0.966 | 0.958 |
|  |  | 2 | **0.982** | 0.958 | 0.978 | 0.951 | 0.800 |
|  |  | 3 | **0.991** | 0.990 | 0.988 | 0.987 | 0.983 |
| RBF kernel SVM | 16 | 1 | **0.976** | **0.976** | 0.974 | 0.973 | 0.956 |
|  |  | 2 | **0.982** | 0.967 | **0.982** | 0.963 | 0.512 |
|  |  | 3 | 0.991 | **0.992** | 0.990 | 0.990 | 0.985 |
|  | 98% | 1 | **0.979** | **0.979** | 0.976 | 0.967 | 0.955 |
|  |  | 2 | **0.984** | 0.970 | 0.983 | 0.966 | 0.531 |
|  |  | 3 | **0.993** | **0.993** | 0.991 | 0.991 | 0.977 |
| random forest | 16 | 1 | **0.970** | **0.970** | 0.967 | 0.965 | 0.943 |
|  |  | 2 | **0.980** | 0.976 | 0.977 | 0.970 | 0.693 |
|  |  | 3 | 0.988 | **0.989** | 0.986 | 0.986 | 0.979 |
|  | 98% | 1 | **0.972** | 0.971 | 0.967 | 0.968 | 0.950 |
|  |  | 2 | **0.979** | 0.971 | 0.975 | 0.965 | 0.696 |
|  |  | 3 | 0.989 | **0.990** | 0.985 | 0.989 | 0.982 |

To assess performance of the different techniques across training set sizes (Fig. 4), we measured the area under the curve for each of the 50 iterations for a given setting. This renders an aggregated performance for each train/test split across the range of training set sizes we used and gives us an estimate of performance for small to medium training set sizes. In contrast to the microaveraged AUC across different training set sizes, this measure takes into account the fact that we tested more smaller training set sizes (in the range of 100-200 samples) and is a more conservative measure than simple averaging in our case. This is achieved by weighting performance according to train set size sampling frequency. Additionally, we normalized performance, so that a perfect classifier would achieve an relative performance of 1, corresponding to an AUC of 1 for all training set sizes in the range from 100 to 400 samples. Fig. 5 shows the distribution of this performance measure for different techniques, classifiers and transfer directions. Across all settings, the *EasyAdapt* technique consistently showed superior performance over other techniques: Among 18 different settings, *EasyAdapt* ranked 15 times the best or tied for the best and 3 times as the second best. This not only demonstrates the effectiveness of knowledge transfer of *EasyAdapt*, but also shows its generality with respect to base classifiers and feature selection methods under different transfer situations. The second best technique was the *Domain* technique, with 8 times the best or tied for the best and 3 times in the second place. This indicated that in many cases the membership feature used by the *Domain* technique was also able to leverage some knowledge from related domains. The technique with the lowest performance was the *Source* technique, which ranked last in every setting.

**Fig. 5.**
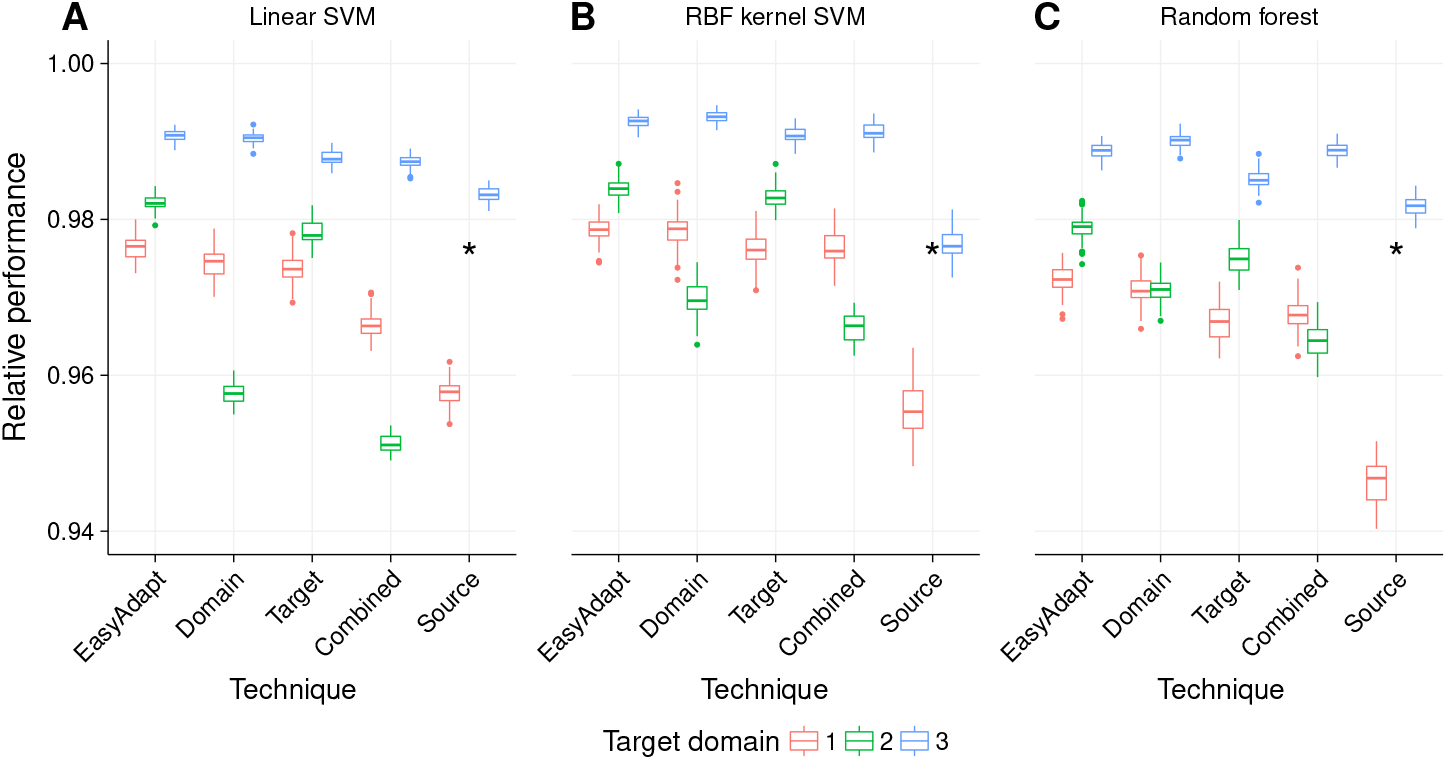
Relative performance, measured as area under the curve for each of the 50 iterations that were used to generate the average performance lines in Fig. 4. Each data point shows performance over the range of training set sizes (100-400) for one iteration of the target domain; each box plot comprises data from 50 iterations. Performance is shown for (A) linear SVM, (B) radial basis function (RBF) kernel SVM, and (C) random forest classifiers.

In practice, it is hard to predict whether pooling of data will actually improve prediction performance or lead to negative transfer, i.e. learning in the target domain might be negatively affected by the use of additional information, if domains are too different [11,27]. An example for such negative transfer is the case of experiment 2 as the target domain. Here, both the *Combined* and *Domain* techniques performed considerably worse compared to the *Target* technique (see Table 1). This can probably be explained by stronger differences in distributions between experiments 1 and 3 on the one hand, and experiment 2 on the other, as experiment 2 used a different magnification. This difference can also be seen from the extremely poor performance of the *Source* technique for experiment 2 as the target domain. It is worth noting that the negative transfer that affected the *Combined* and *Domain* techniques with experiment 2 as target domain appears stable across different training set sizes (Fig. 4). Importantly, we do not observe such negative transfer in the case of the *EasyAdapt* technique. Performance of *EasyAdapt* was comparable or even slightly better than the *Target* technique when looking at experiment 2 as the target domain.

## 4 Discussion

In the present study, we investigated whether accounting for experimental variation in biological data using a domain adaptation techniques can help improve prediction performance and reduce the need for labeled data. We show that indeed, given only limited training data, the *EasyAdapt* domain adaptation technique boosts prediction performance both in a simulation study and a data set of imaged single cells [25] and leads to more robust predictions in the presence of experimental variation.

Recently, there have been a number of approaches that try to improve generalization of deep neural network performance across multiple domains. This is important, as neural networks have been known to generalize relatively poorly [28]. Often, the approach is to learn transferable representations that both identify the factors driving variation within the data and match feature distributions across domains [29,30]. Recent work has used models that are able to adapt to different domain very quickly by using an efficient parametrization of deep neural networks and adapter residual modules [31,32]. There is also interesting work combining generative adversarial networks with domain adaptation [33,34,35]. It is worth noting that the approach described in this work is orthogonal to these models, and can be used with any type of supervised machine learning algorithm, including but not limited to deep neural networks.

Applications of domain adaptation techniques in biological research have so far been mostly restricted to genomic sequence analysis [36,37]. Widmer et al. [38,39] used a more general multi-task learning framework in conjunction with regularization based supervised learning methods, such as SVM and logistic regression for splice-site and binding site prediction and to transfer model parameters learned on 2D images to 3D images in order to enhance learning. In contrast to [39], we do not learn domain specific differences explicitly. In practice, this information is also often hard to quantify. Here, we rather focus on the effect of training set size and the pooling of heterogeneous data without quantitative knowledge about the relationship between domains. We compare performance of the *EasyAdapt* technique across three different machine learning algorithms. Furthermore, we consider a range of common ways of combining information from different domains, e.g. via explicit encoding of domain membership, a procedure that is often used in practice. We demonstrate that the *EasyAdapt* technique is relatively robust to negative effects of data pooling.

Our results have implications for dealing with biological batch effects in machine learning tasks and for improving learning in settings with limited training data, if additional source data is available. The *EasyAdapt* technique allows the reuse of existing data sets as source data and avoids cost-intensive manual labelling of training data. Results confirm the problem that is one major motivation of this work: a model trained using data from one biological experiment is likely to have much inferior performance when applied to a different experiment, despite the experiments sharing similar experimental setups. Importantly, the *EasyAdapt* technique is general in that it does not change the machine learning method used and can therefore be applied to a wide set of problems. Because the feature space grows linearly in the number of domains, the approach is not applicable in cases with very large feature spaces or a large number of domains.

In general, classification accuracy in the transfer learning setting will be an increasing function of both the number of training samples available and the homogeneity and level of relatedness of the training samples to the test set. Given a limited set of training samples and reasonable relatedness between training and test set, transfer learning can help to improve classification accuracy. However, in the case when the relatedness between training and test set is insufficient to enable transfer, there is potential for negative impact when adding additional data from a different domain (known as negative transfer). *EasyAdapt* strikes a balance between improving performance in cases when additional information is available and robustness to experimental variations. Compared with classic techniques such as the *Domain* and *Combined* techniques, the *EasyAdapt* technique is less affected by negative transfer and for small to medium training set sizes it can improve learning in the target domain.

The technique is limited by the necessity to identify domains, i.e. it is necessary to have domain knowledge about potential differences in experimental conditions and fundamental differences in feature distributions that define domains. Furthermore, it requires that the domains have a shared feature subspace and are distinct [16]. Both requirements are typically fulfilled in biological data. Further research will be necessary to develop empirical measures of domain relationships that help to identify cases where the use of domain adaptation in machine learning can be particularly helpful.

## Notes

#### Summary of Updates

Slightly revised and accepted at the 28th International Conference on Artificial Neural Networks (ICANN) 2019 in Munich, Germany

## References

1. Golkov, V., Skwark, M.J., Golkov, A., Dosovitskiy, A., Brox, T., Meiler, J., Cremers, D.: Protein contact prediction from amino acid co-evolution using convolutional networks for graph-valued images. In: Advances in Neural Information Processing Systems. pp. 4222–4230 (2016)

2. Rost, B., Sander, C.: Combining evolutionary information and neural networks to predict protein secondary structure. Proteins: Structure, Function, and Bioinformatics 19(1), 55–72 (1994). https://doi.org/10.1002/prot.340190108

3. Xiong, H.Y., Alipanahi, B., Lee, L.J., Bretschneider, H., Merico, D., Yuen, R.K.C., Hua, Y., Gueroussov, S., Najafabadi, H.S., Hughes, T.R., Morris, Q., Barash, Y., Krainer, A.R., Jojic, N., Scherer, S.W., Blencowe, B.J., Frey, B.J.: The human splicing code reveals new insights into the genetic determinants of disease. Science 347(6218), 1254806 (Jan 2015). https://doi.org/10.1126/science.1254806

4. Alizadeh, A.A., Eisen, M.B., Davis, R.E., Ma, C., Lossos, I.S., Rosenwald, A., Boldrick, J.C., Sabet, H., Tran, T., Yu, X., Powell, J.I., Yang, L., Marti, G.E., Moore, T., Hudson, J., Lu, L., Lewis, D.B., Tibshirani, R., Sherlock, G., Chan, W.C., Greiner, T.C., Weisenburger, D.D., Armitage, J.O., Warnke, R., Levy, R., Wilson, W., Grever, M.R., Byrd, J.C., Botstein, D., Brown, P.O., Staudt, L.M.: Distinct types of diffuse large B-cell lymphoma identified by gene expression profiling. Nature 403(6769), 503–511 (2000). https://doi.org/10.1038/35000501

5. Helmstaedter, M., Briggman, K.L., Turaga, S.C., Jain, V., Seung, H.S., Denk, W.: Connectomic reconstruction of the inner plexiform layer in the mouse retina. Nature 500(7461), 168–174 (2013)

6. Buggenthin, F., Buettner, F., Hoppe, P.S., Endele, M., Kroiss, M., Strasser, M., Schwarzfischer, M., Loeffler, D., Kokkaliaris, K.D., Hilsenbeck, O., et al.: Prospective identification of hematopoietic lineage choice by deep learning. Nature methods 14(4), 403 (2017)

7. Blasi, T., Hennig, H., Summers, H.D., Theis, F.J., Cerveira, J., Patterson, J.O., Davies, D., Filby, A., Carpenter, A.E., Rees, P.: Label-free cell cycle analysis for high-throughput imaging flow cytometry. Nature Communications 7, 10256 (2016). https://doi.org/10.1038/ncomms10256

8. Leek, J.T., Scharpf, R.B., Bravo, H.C., Simcha, D., Langmead, B., Johnson, W.E., Geman, D., Baggerly, K., Irizarry, R.A.: Tackling the widespread and critical impact of batch effects in high-throughput data. Nature Reviews Genetics 11(10), 733 (2010)

9. Bernau, C., Riester, M., Boulesteix, A.L., Parmigiani, G., Huttenhower, C., Waldron, L., Trippa, L.: Cross-study validation for the assessment of prediction algorithms. Bioinformatics 30(12), i105–i112 (2014)

10. Patricia, N., Caputo, B.: Learning to learn, from transfer learning to domain adaptation: A unifying perspective. In: Proceedings of the IEEE Conference on Computer Vision and Pattern Recognition. pp. 1442–1449 (2014)

11. Pan, S.J., Yang, Q.: A survey on transfer learning. IEEE Transactions on knowledge and data engineering 22(10), 1345–1359 (2010)

12. Patel, V.M., Gopalan, R., Li, R., Chellappa, R.: Visual Domain Adaptation: A survey of recent advances. IEEE Signal Processing Magazine 32(3), 53–69 (2015). https://doi.org/10.1109/MSP.2014.2347059

13. Hwa, R.: Supervised Grammar Induction Using Training Data with Limited Constituent Information. In: Proceedings of the 37th Annual Meeting of the Association for Computational Linguistics on Computational Linguistics. pp. 73–79. Association for Computational Linguistics, Stroudsburg, PA, USA (1999). https://doi.org/10.3115/1034678.1034699

14. Gildea, D.: Corpus variation and parser performance. In: Proceedings of the 2001 Conference on Empirical Methods in Natural Language Processing. pp. 167–202 (2001)

15. Daume III, H., Marcu, D.: Domain adaptation for statistical classifiers. Journal of artificial Intelligence research 26, 101–126 (2006)

16. Daumé III, H.: Frustratingly Easy Domain Adaptation. ACL p. 256 (2007)

17. Laing, E.E., Möller-Levet, C.S., Poh, N., Santhi, N., Archer, S.N., Dijk, D.J.: Blood transcriptome based biomarkers for human circadian phase. eLife 6, e20214 (2017). https://doi.org/10.7554/eLife.20214

18. Chen, L., Qu, X., Cao, M., Zhou, Y., Li, W., Liang, B., Li, W., He, W., Feng, C., Jia, X., He, Y.: Identification of breast cancer patients based on human signaling network motifs. Scientific Reports 3 (2013). https://doi.org/10.1038/srep03368

19. Wang, X., Naqa, E., M, I.: Prediction of both conserved and nonconserved microRNA targets in animals. Bioinformatics 24(3), 325–332 (2008). https://doi.org/10.1093/bioinformatics/btm595

20. Hsu, C.W., Chang, C.C., Lin, C.J., et al.: A practical guide to support vector classification (2003)

21. Daumé, III, H., Kumar, A., Saha, A.: Frustratingly Easy Semi-supervised Domain Adaptation. In: Proceedings of the 2010 Workshop on Domain Adaptation for Natural Language Processing. pp. 53–59. Association for Computational Linguistics (2010)

22. Cortes, C., Vapnik, V.: Support-vector networks. Machine Learning 20(3), 273–297 (1995). https://doi.org/10.1007/BF00994018

23. Boser, B.E., Guyon, I.M., Vapnik, V.N.: A training algorithm for optimal margin classifiers. In: Proceedings of the fifth annual workshop on Computational learning theory. pp. 144–152. ACM (1992)

24. Sokolova, M., Lapalme, G.: A systematic analysis of performance measures for classification tasks. Information Processing & Management 45(4), 427–437 (2009). https://doi.org/10.1016/j.ipm.2009.03.002

25. Held, M., Schmitz, M.H.A., Fischer, B., Walter, T., Neumann, B., Olma, M.H., Peter, M., Ellenberg, J., Gerlich, D.W.: CellCognition: time-resolved phenotype annotation in high-throughput live cell imaging. Nature Methods 7(9), 747–754 (2010). https://doi.org/10.1038/nmeth.1486

26. Breiman, L.: Random Forests. Machine Learning 45(1), 5–32 (2001). https://doi.org/10.1023/A:1010933404324

27. Rosenstein, M.T., Marx, Z., Kaelbling, L.P., Dietterich, T.G.: To transfer or not to transfer. In: NIPS 2005 workshop on transfer learning. vol. 898, pp. 1–4 (2005)

28. Yosinski, J., Clune, J., Bengio, Y., Lipson, H.: How transferable are features in deep neural networks? In: Advances in Neural Information Processing Systems. pp. 3320–3328 (2014)

29. Ganin, Y., Ustinova, E., Ajakan, H., Germain, P., Larochelle, H., Laviolette, F., Marchand, M., Lempitsky, V.: Domain-Adversarial Training of Neural Networks. In: Csurka, G. (ed.) Domain Adaptation in Computer Vision Applications, pp. 189–209. Springer International Publishing, Cham (2017)

30. Long, M., Zhu, H., Wang, J., Jordan, M.I.: Deep Transfer Learning with Joint Adaptation Networks. In: Proceedings of the 34th International Conference on Machine Learning – Volume 70. pp. 2208–2217 (2017)

31. Rebuffi, S.A., Bilen, H., Vedaldi, A.: Learning multiple visual domains with residual adapters. In: Advances in Neural Information Processing Systems 30, pp. 506–516 (2017)

32. Rebuffi, S.A., Bilen, H., Vedaldi, A.: Efficient Parametrization of Multi-Domain Deep Neural Networks. In: Proceedings of the IEEE Conference on Computer Vision and Pattern Recognition. pp. 8119–8127 (2018)

33. Tzeng, E., Hoffman, J., Darrell, T., Saenko, K.: Simultaneous Deep Transfer Across Domains and Tasks. In: Proceedings of the IEEE International Conference on Computer Vision. pp. 4068–4076 (2015)

34. Tzeng, E., Hoffman, J., Saenko, K., Darrell, T.: Adversarial Discriminative Domain Adaptation. In: Proceedings of the IEEE Conference on Computer Vision and Pattern Recognition. pp. 7167–7176 (2017)

35. Long, M., Cao, Z., Wang, J., Jordan, M.I.: Conditional Adversarial Domain Adaptation. In: Advances in Neural Information Processing Systems 31, pp. 1640–1650 (2018)

36. Jacob, L., Vert, J.P.: Efficient peptide–mhc-i binding prediction for alleles with few known binders. Bioinformatics 24(3), 358–366 (2007)

37. Schweikert, G., Rätsch, G., Widmer, C., Schölkopf, B.: An empirical analysis of domain adaptation algorithms for genomic sequence analysis. In: Advances in Neural Information Processing Systems. pp. 1433–1440 (2009)

38. Widmer, C., Rätsch, G.: Multitask learning in computational biology. In: Proceedings of ICML Workshop on Unsupervised and Transfer Learning. pp. 207–216 (2012)

39. Widmer, C., Kloft, M., Lou, X., Rätsch, G.: Regularization-based multitask learning with applications to genome biology and biological imaging. KI-Künstliche Intelligenz 28(1), 29–33 (2014)

